# Detection of rare disease-related genetic variants using the birthday model

**DOI:** 10.1101/464842

**Authors:** Yael Berstein, Shane E. McCarthy, Melissa Kramer, W. Richard McCombie

**Affiliations:** The Stanley Institute for Cognitive Genomics, Cold Spring Harbor Laboratory, Cold Spring Harbor, New York, USA

## Abstract

**Motivation:** Exome sequencing is a powerful technique for the identification of disease-causing genes. A number of Mendelian inherited disease genes have been identified through this method. However, it remains a challenge to leverage exome sequencing for the study of complex disorders, such as schizophrenia and bipolar disorder, due to the genetic and phenotypic heterogeneity of these disorders. Although not feasible for many studies, sequencing large sample sizes (>10,000) may improve statistical power to associate more variants, while the aggregation of distinct rare variants associated with a given disease can make the identification of causal genes statistically challenging. Therefore, new methods for rare variant association are imperative to identify causative genes of complex disorders.

**Results:** Here we propose a method to predict causative rare variants using a popular probabilistic problem: The Birthday Model, which estimates the probability that multiple individuals in a group share the same birthday. We consider the probability and coincidence of samples sharing a variant akin to the chance of individuals sharing the same birthday. We investigated the parameter effects of our model, providing guidelines for its use and interpretation of the results. Using published data on autism spectrum disorder, hypertriglyceridemia in addition to a current case-control study on bipolar disorder, we evaluated this probabilistic method to identify potential causative variants. Several genes in the top results of the case-control study were associated with autism spectrum and bipolar disorder. Given that the core probability based on the birthday model is very sensitive to low recurrence, the method successfully tests the association of rare variants, which generally do not provide enough signal in commonly used statistical tests. Importantly, the simplicity of the model allows quick interpretation of genomic data, enabling users to select gene candidates for further biological validation of specific mutations and downstream functional or other studies.

**Availability:** https://github.com/yberstein/Birthday-Alqorithm http://labshare.cshl.edu/shares/mccombielab/www-data/Birthday-Algorithm/Birthday-Alqorithm.html

**Contact:** (or)

**Supplementary information:** Supplementary data are available online.

## 1 Introduction

Next generation sequencing (NGS), both exome and whole genome (Goodwin, et al., 2016; Shendure, et al., 2017), is a powerful tool for the investigation of genomic sequences and the identification of disease-related genes (Boycott, et al., 2013; Chong, et al., 2015; Do, et al., 2015; Stranneheim and Wedell, 2016; Yoshida, et al., 2011). It poses an exciting opportunity to investigate thousands of diseases for which causative genes remain to be discovered. Notably, complex diseases such as schizophrenia, bipolar disorder, autism and major depression, are particularly hard to study due to their genetic and phenotypic heterogeneity (Agarwala, et al., 2013; Hindorff, et al., 2011; McClellan and King, 2010; Mitchell, 2012; Mitchell and Porteous, 2011; Pritchard and Cox, 2002). Hundreds of genetic loci associated with complex disorders have been detected by genome-wide association studies (GWAS). However, the vast majority of these peaks have not been linked to a specific coding or non-coding variant that contributes to the disease and they can only explain a small portion of the expected heritability of these diseases (Eichler, et al., 2010; Maher, 2008; Manolio, et al., 2009). This discrepancy between phenotypic and genetic evidence, often referred to as “missing heritability”, is a key barrier to understanding complex disorders.

A working hypothesis in the field of psychiatric genetics is the idea that certain diseases can be caused by many different variants in many different genes, leading to mechanistic complexity. As a result, case-control genomic studies using standard metrics of statistical significance derive multiple and widespread low frequency signals. Assigning likelihood probabilities to subtle variants of this type remains a major obstacle for the identification of causative variants in complex disorders. A simulation-based study (Kryukov, et al., 2009) suggested that at least 10,000 individuals are required to ensure satisfactory statistical power for case-control genomic studies. Another analytical study (Zuk, et al., 2014) recommends at least 25,000 cases together with a replication cohort. Accordingly, similar recommendations were proposed to overcome relatively low frequencies observed in putative causative variants (Kiezun, et al., 2012; Tennessen, et al., 2012; Zollner, 2012). Finally, a comparison of methods to detect disease-associated variants showed that even sample sizes of ten thousand individuals made little improvement in statistical power, further suggesting that much larger sample sizes are needed (Moutsianas, et al., 2015). In light of these observations, the cost and logistic complexity of genomic studies could significantly escalate, despite the dramatic drop in sequencing costs in recent years.

In this work, we introduce a predictive method to successfully identify possible causal genes and variants in smaller sample sizes in complex disorders. We assign the probabilistic likelihood of rare variants by assessing the probability that a recurrent mutation within a gene of a given coding length would happen by chance in a group of individuals of a given size. To this aim, we represented genomic data as a popular probabilistic problem, the birthday problem, which estimates the probability that a group of individuals share the same birthday. Our solution provides intuitive and simple criteria to prioritize gene variant candidates from genomic data for further downstream investigation. Here, we will show implementations on exome sequencing analysis.

## 2 Materials and Methods

The predictive method for prioritizing rare variants incorporates two main elements. The core element is the probability that a variant is observed by chance in a group of individuals; this is derived from the famous birthday probability. The second element is the way we implement this core probability in order to evaluate each observed variant. This is the decisionmaking method, which combines both the probabilities in the group of cases and in the group of controls to the decision process. Finally, the algorithm implements a permutation method to evaluate all variants observed in the analysis, and provides the final ranking of potentially causing rare variants.

### 2.1 The birthday probability

We simplify the problem of rare variant association by focusing on the probability that individuals would have the same variant by chance. The analogy to the birthday problem is straightforward: in a group of N individuals, what is the probability that at least k of them share the same birthday by chance or analogously, have the same mutated variant? The formulation is based on the following model.

#### Probability model

Adopting the generalized birthday model, first presented by Mckinney (Mckinney, 1966), and its approximated solution (Diaconis and Mosteller, 1989), the probability of coincidence *p* can be extracted from

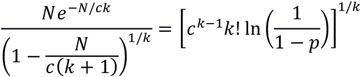

where *N* is the sample size, *c* is the coding length of the gene where the variant was observed and k is the observed recurrence of this variant i.e. the number of individuals presenting this same variant.

### 2.2 The decision-making method

In a cohort of exome sequenced cases and controls, we observe for each SNP (single-nucleotide polymorphism) the total number of cases and the total number of controls that present this specific variant. Then, we question if our observation is a mere coincidence or if it may be indicating a potential causative variant. Ascertaining the probability of coincidence in both cases and controls allow us to evaluate our observation. If the probability results in a high value, it indicates that this observation is probably occurring by chance. Conversely, an indication of a real finding is when the probability results in a low value, indicating that there is a low probability of the observation being a coincidence. Generally, we expect that a causative variant would be observed in cases, but not in controls. Therefore, the decision-making method selects variants that simultaneously show low probability of coincidence in cases and high probability of coincidence in controls. The birthday problem probability is quite sensitive to absolute values of recurrence and sample size. Therefore, even in the case of rare variants, where recurrence is low, this model should be sensitive to small deviations from the recurrence rates expected for coincidental events.

### 2.3 The algorithm

The user could define the accepted level of coincidence; however, this would lead to a subjective decision-making based on the specific threshold. For this reason, we propose the following multiple testing algorithm inspired by Westfall & Young (Westfall and Young, 1993). First, the algorithm computes for each variant the probability of the observed recurrence in cases and controls, *p*_case_* and *p*_control_* respectively. Then, after permuting the disease status of the original dataset several times, it computes for each variant the probability in cases and controls for each permutated dataset. The final probability of each variant is the proportion of permutations where both the probability of cases and controls were more extreme than the observed probabilities. The permutations that will be counted are those that simultaneously have a lower probability in cases than the observed in the sample of cases and higher probability in controls than the observed probability in the sample of controls. Importantly, the algorithm can be applied at both variant and gene resolution. In a gene level implementation, the count for recurrence within a gene is defined as the overall recurrence of all variants in the specific gene.

#### Algorithm

1. Compute *p*_case_* and *p*\*_control_ for each gene/variant.
2. Permutation of the disease status of the individuals.
3. For each permutation compute *p_case_* and *p_control_* for each gene/variant.
4. Compute for each gene/variant:

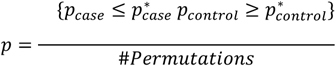
5. Rank the genes/variant in increasing order of p.

## 3 Results

We tested several aspects of the proposed method in both simulated data and in published data. The simulated data consists of three different sets. In simulation set I, we verified that our adaptation of the birthday model could properly represent the probability of recurrence of a variant. We have tested the behavior of the core probability under different parameters. Second, in simulation set II, we tested the ability of the birthday probability to test the association of rare variants to a disease in several simulated case-control studies. In the last set of simulations, set III, we tested the limit of the model in an actual case-control study of bipolar disorder (Goes, et al., 2016) using a wide range of recurrence values. We then tested the decision-making method on two published data sets of: autism spectrum disorder (Iossifov, et al., 2014) and hypertriglyceridemia (Johansen, et al., 2010). The implementation of our model on these two studies is based on our initial method which did not employ a permutation step. Therefore, the analysis relies on the comparison of the resulting probability of cases and of controls, which gave valuable insight about the basic model. Finally, we fully implemented our algorithm in a case-control study on bipolar disorder (Goes, et al., 2016) with ten thousand permutations. Here we compared the method with PLINK/SEQ and SKAT, which were the methods used in Goes study.

### 3.1 Simulations

#### 3.1.1 General behavior of the core probability - simulation set I

The generalized birthday model provides an estimate of the probability that a recurrent mutation occurred by random chance. Changes in each one of the parameters of the model – sample size (*N*), gene size (*c*) and recurrence (*k*) - lead to changes in the probabilities as expected, as illustrated in figure 1. In this simulation set we computed the core probability based on the equation (Diaconis and Mosteller, 1989) described in section 2.1 and considering the following values for the parameters:

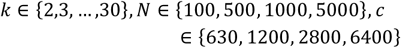

**Fig. 1.**
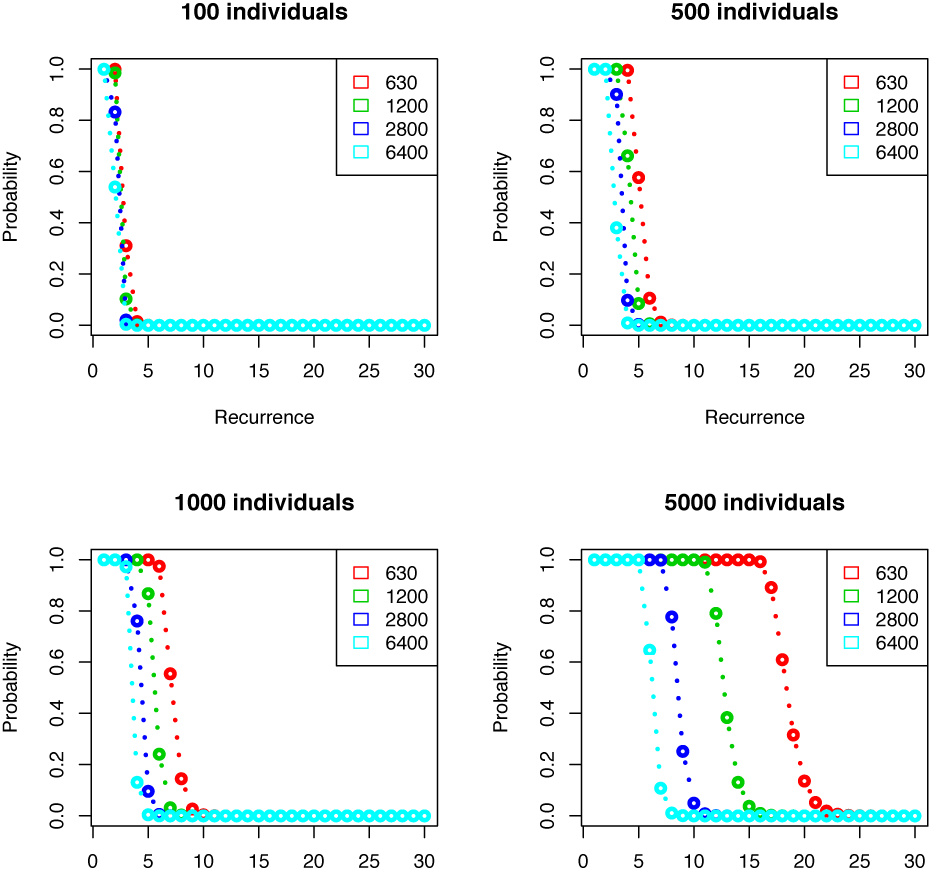
Impact of gene coding length and recurrence rates on the Birthday Method for variant detection. The plots show the probabilities (vertical axis) computed by the birthday model at different levels of observed recurrence (horizontal axis). Each curve represents a gene with a different coding length: 630 nucleotides represents the coding length of 20% of the genes, 1200 nn 50%, 2800 nn 90% and 6400 nn 99% (based on principal isoform, APPRIS database). The distribution of the coding length based on Appris database can be found in supplementary fig. 1

The coding length of a gene used in this simulation, represented by c, is based on the distribution of the principal isoform length (Rodriguez, et al., 2013). Specifically, 630 nucleotides represent the coding length of 20% of the genes, 1200 nn 50%, 2800 nn 90% and 6400 nn 99%. More details on the coding size of the gene can be found in supplementary material.

We have tested the probability of coincident recurrence computed by our model while varying only one of the three parameters. (i) We observe that for fixed sample size and recurrence level, as the coding length of a gene increases the probability of coincident recurrence decreases, i.e. it would be less likely for a variant to coincide on the same nucleotide by chance as gene length increases. (ii) When testing the probability of coincidence in genes of the same coding length and recurrence, the probability of coincidence increases as the sample size becomes larger. i.e. more individuals increase the chances of coincidences happening. (iii) Observing the negative slope of all curves in figure 1, we see that for the same sample size and same coding length as the recurrence rates increase the probability of coincident recurrence decreases, which is what we would expect since higher recurrence of a mutation may indicate an interesting observation.

#### 3.1.2 Performance of the decision-making method in rare variant association – simulation set II

The following set of simulations illustrates the ability of the decision-making model to identify rare variants associated with a disease. Each instance of the simulation is defined by three parameters: the sample size(*N*), the recurrence of the implanted causative variant in cases, and the recurrence of the implanted causative variant in controls. For simplicity, the sample size is the same in cases and controls and varied from 50, 500 and 5000 individuals. Given that our focus is on the identification of rare variants associated with a disease, we set the recurrence of the implanted causative variant to be 5 or 10 in cases, and 1, 2, 3 or 4 in controls. Note that the fraction of the sample population sharing a given variant is determined by both the recurrence and the sample size. For example, a fraction of 20% is derived from recurrence of 10 in a sample of 50 individuals (10/50), and it represents a relatively common variant shared among 20% of the sample population. Whereas the same recurrence of 10 individuals in a sample of 5000 defines a fraction of 0.2% (10/5000), and represents a rare variant. In order to evaluate the limit of the model, we ran 1000 iterations of each simulation instance. The recurrence of the background variant was simulated based on allele frequencies of the Exome Variant Server, NHLBI GO Exome Sequencing Project (ESP), Seattle, WA (URL: http://evs.gs.washington.edu/EVS/), for details see supplementary figure 2. The coding size of each gene is sampled from the principal isoform distribution in APPRIS database. For each instance, we calculated the number of iterations where the implanted causative variant was the top 1 (red), top 2 (green) or top 3 (blue) variant based on the basic probability method and 5% accepted level of coincidence (Figure 2). We observe that for sample sizes of 50 and 500 individuals there are several instances where the implanted causative variant was ranked at the top in almost all of the 1000 iterations (Figure 2). For example, even when sample size is 500 individuals, a variant recurring in 2% (10/500) of cases and 0.08% (4/500) in controls can be detected. As the sample size increases, the relative fraction of patients sharing a given variant decreases (representing rarer variants), therefore is expected to have a lower success in detecting the implanted causative variant as illustrated in figure 2.

**Fig. 2.**
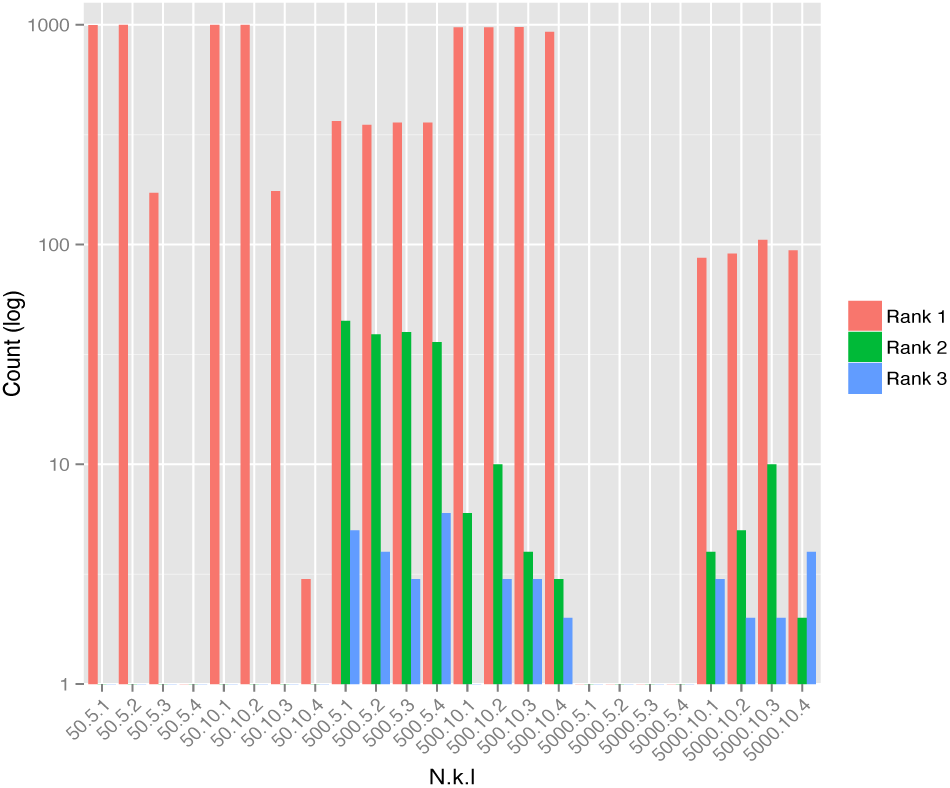
In the horizontal axis are scenarios defined by N-sample size, k – recurrence in cases and 1 – recurrence in control. Red bar represents the number of iterations that the implanted “causative” variant was ranked in the first place, green second place and blue third.

#### 3.1.3 Testing the limits of the decision-making method in a bipolar disorder study – simulation set III

Here we evaluate the decision-making method for a broader range of recurrence of the implanted causative variant. We focus on simulations of a case-control study on bipolar disorder (Goes, et al., 2016), comprised of 1141 affected individuals and 1146 unaffected controls, as a representation of a study in a complex disorder. The recurrence of each variant in cases and controls, and the coding length of this simulation set are generated similarly to the previous simulation set. In addition to that, we simulated different “causative” rare variant scenarios by implanting a “causative” variant such that k, 1 ∈ {1,2, …, 20 |k>l}. For each instance, we ran 1000 iterations. Figure 3 illustrates the proportion of iterations in each instance for which the implanted causative variant was ranked in the top 3 based on the method. We observe that already with recurrence in cases (k) of 11 individuals, which is around 1% of cases, the model has an ~90% of success even when there are 5 controls with the same mutation. Note that even in a scenario of 6 cases having the same causative variant, the model detects the causative mutation in about 20% of the iterations.

**Fig. 3.**
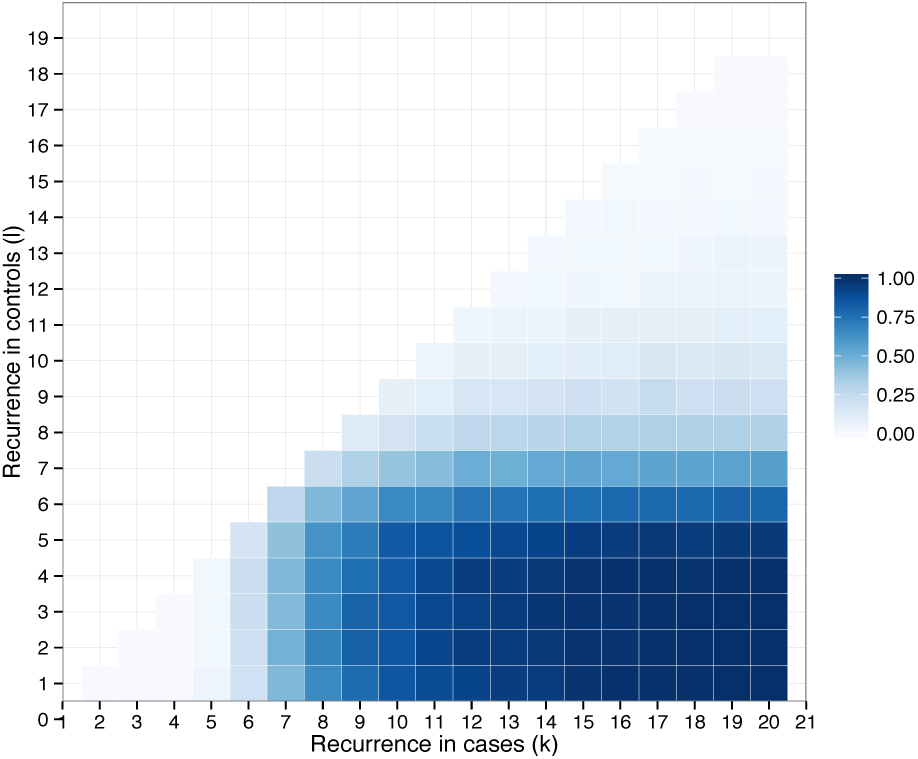
Each tile in this heat map represents a different scenario defined by the recurrence in cases (k, horizontal axis) and in controls (1, vertical axis) of the implanted “causative” variant. The sample sizes are fixed to 1141 cases and 1146 controls, as in the bipolar study. The color shade represents the proportion on iterations that our method detected the “causative” variant and ranked it in first three places. Darker blue indicates a higher proportion of success.

### 3.2 Implementation of the decision-making method on real data

#### 3.2.1 Autism spectrum disorder

Here we illustrate the adaptability of the model to different studies. We show an implementation of the decision-making method to a family study design, as opposed to the previously described case-control study implementation. We used the birthday model on data from a study on autism-spectrum disorder (Iossifov, et al., 2014). Even though the method was developed for case control studies, we can implement it on family studies that focus on de novo mutations, as in the study on autism, as follows: de novo mutations are those detected in the affected sibling but not in either of the parents. The autism study consisted of exome sequencing to identify de novo mutations in simplex families: trios (unaffected parents with one affected child) or quads (unaffected parents with one affected and one unaffected child). Given that the families are independent, in order to implement our decision-making method, we group the affected children as the case samples and the unaffected children as the control samples. The study focused on “likely gene disruptive” mutations (LGD) which include frameshift, splice-site and nonsense mutations. Therefore, the input for recurrence is restricted to the counts of LGD mutations in cases and controls in each gene. Specifically, 2508 affected children and 1911 unaffected siblings. According to our model, only gene CHD8 presents a relatively low probability of coincidence in cases (0.043%), and simultaneously high probability in controls (100%). Therefore, this is the only gene that the birthday model would indicate as worthy of further investigation. Note that the original study (Iossifov, et al., 2014) reported significant results for specific gene-sets, such as chromatin modifier and FMRP target genes. However, in a gene level analysis, no specific gene had a statistically significant number of de novo mutations in cases compared to controls, even CHD8 which was the most recurrently hit gene. We observe that all other genes showed a very high probability of being a coincidence in cases, as described in Supplementary Table 1. These results show the behavior of the model when relatively low recurrence is observed in the case group. When applying the model to LGD mutations, the model predicts only CHD8 to be a potential candidate. We also observe that none of the variants were recurrent in this gene, meaning that there were 7 different variants detected. Therefore, a variant level implementation would not select any of the variants of this gene. We also implemented the model after collapsing the counts of LGD and missense mutations, even though they are separate categories in the original paper, and we observed that gene SCN2A has 6 mutations in affected individuals (2 LGD and 4 missense), therefore 2.82% probability of being a coincident recurrently hit gene in 6 different individuals.

**Table 1.**
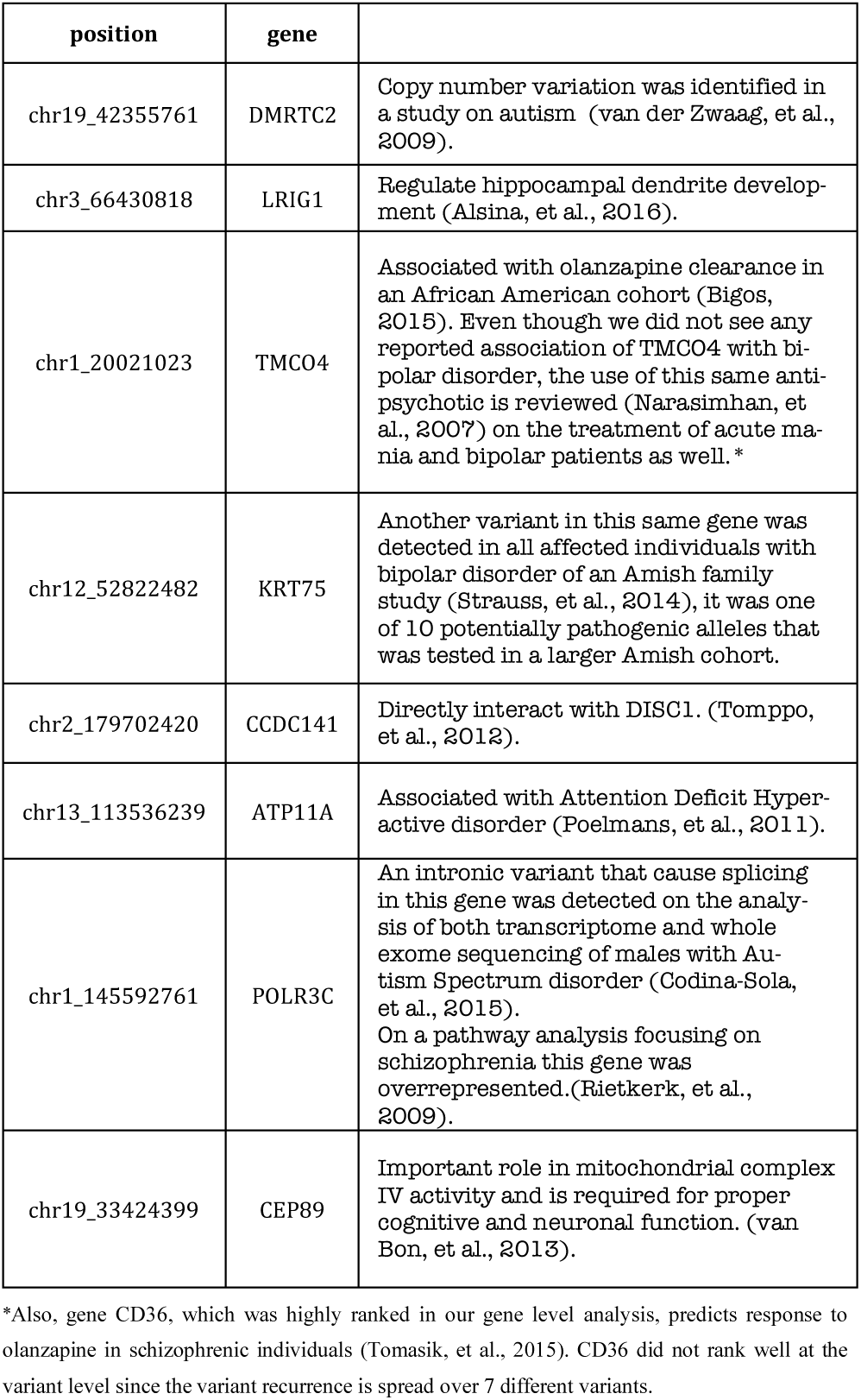
This table describes the supporting findings of the top 15 variants which are located in genes already associated with psychiatric disorders.

#### 3.2.2 Hypertriglyceridemia

We implemented the decision-making method based on the birthday model on a study of hypertriglyceridemia (Johansen, et al., 2010). We analyzed this data at both the variant level and gene level, as an illustration of the differences of the decision-making method when applied at different resolutions. First, they did a GWAS study to identify common variants associated with hypertriglyceridemia. Then, they resequenced the genes that were identified in the first stage in order to identify rare variants within these genes. Because our focus is on rare variant association, we concentrate on this second part of the study. It is comprised of four genes, APOA5, GCKR LPL and APOB (exons 26 and 29, 67.8% of the coding region of this gene) that were re-sequenced in 438 cases and 327 controls. There were 80 rare variants, with minor allele frequency of at most 1% in controls.

We first performed a variant level analysis based on the birthday model. Supplementary Table 2 describes for each one of the 80 rare variants, its respective recurrence in cases and in controls, and the resulting probability of cases and of controls. The model prioritizes 6 variants that presented very low probability in cases and very high probability in controls: p.Q234P in gene GCKR, p.G188E in gene LPL, and p.E2539K, p.P2794L, and p.S3252G in gene APOB. No variant in gene APOA5 showed interesting probabilities, as all variants presented probability value 1 in both cases and controls. In this data set the probability values were quite extreme i.e. values close to zero or one in both cases and controls. Therefore, regardless of the cutoff level, the same set of variants would be selected. We observe the expected: the decision model will “exclude” the rare variant that recurred only one time in a case or in both groups.

Then, we implemented our model at a gene level resolution, using 5% as the threshold of the probability of coincidence, to be consistent with the methods of the authors, who applied a fisher exact test with nominal statistical significance defined as two-sided p<5%. It is important to note that the published study counts the total alleles, thus sample size in the birthday model is equal to twice the number of individuals with diploid genomes. Their analysis combined all variants of the 4 candidate genes showing enrichment of rare variants when comparing the counts in cases versus controls. When performing a gene level analysis, we observe that our decision-making method would not indicate any of these four genes as potential candidates. None of the genes presented simultaneously a relatively low probability in cases and high probability in controls, as described in Supplementary Table 3. Even though, genes GCKR, LPL and APOB have very low probability in cases, the method discarded those genes due to low probabilities in controls as well. In fact, collapsing the variants to the gene level had diluted the signal of these genes due the different variants found in the controls. If sample size were the total number of individuals, as we propose in our method, the results are very similar. It is worth noting that the majority of the individuals carry a single rare variant. This is helpful in our model when applying at a gene level, since we know we were not over-counting recurrence of individuals in the same gene. Also, reported in that article, the few individuals with multiple rare variants were overrepresented among individuals with HTG. Therefore, our model could be inflating the recurrence, leading to a better (lower) probability. Despite this, we could not detect “interesting” genes in the gene level analysis.

### 3.3 Implementation of the algorithm to a study on bipolar disorder

Here we implemented the algorithm based on the birthday model on a recent case-control study on bipolar disorder (Goes, et al., 2016), consisting of 1135 cases and 1142 controls. Note that for implementing the algorithm, which includes the permutation step, the detailed SNP profile of each individual is needed as input. In our implementation, we focus on the set of disruptive variants – nonsense, splice-sites and frameshift mutations, with minor allele frequency of at most 1% in EVS, 1000 Genomes, and the case-control group. Both variant level and gene level analysis were performed, using 10000 permutated datasets. We also compare the results of our algorithm with the methods applied in the original study. At the variant level we compare our results to Firth’s penalized logistic regression (PLR)(Firth, 1993). At the gene level we compare our results with both SKAT (Wu, et al., 2011) and the burden test as implemented in PLINK/SEQ. The variant list in Supplementary Table 4 describes the top 20 ranked variants by our method, along with the rank according to PLR and the recurrence of the specific variant in the case group and in the control group. It also shows the rank of the respective gene from the gene level analysis by our method, by SKAT and by PLINK/SEQ, with the respective recurrence. Note that both our algorithm in variant level resolution and PLR ranked the same top 1 variant in gene ZNF677, but quite different ranking for most of the top 20 variants. At the gene level, the ranking of the top 20 genes by our algorithm is closer to the ranking of SKAT than PLINK/SEQ. We observe overlap between the gene level and the variant level analyses based on our algorithm, especially on the top of the list. For example, the top three variants in the variant level analysis are located in the top three genes of the gene level analysis: ZNF677, DMRTC2 and LY75/LY75-CD302 (the variant is located in a region that overlap these two last). This shows the ability to detect rare variants even when applying the method at “high resolution” (variant level). The consistent results at the gene level resolution reinforce the findings, showing that even at a lower resolution the results are the same. Those top variants and genes are of special interest for follow up. Furthermore, any variants that ranked high, even if the gene in which they are located had a lower ranking should be further investigated. There are several reasons for that, but the main factor is likely to be that the gene level analysis could dilute the signal. As we observed in the previous section, the gene could get ranked low due recurrence of other variants in the same gene in controls. In figure 4, we can see a graphical representation of the recurrence in cases and controls of the top 50 variant by our method. Note that the small difference between the recurrence values, for example 5 cases and zero controls among the top 10 variants, can be detected by our algorithm. In figure 5 we can see the difference of the ranking of our algorithm in variant level compared to PLR, we observe concordance of the algorithms in the top 1 and 3 variants.

**Fig. 4.**
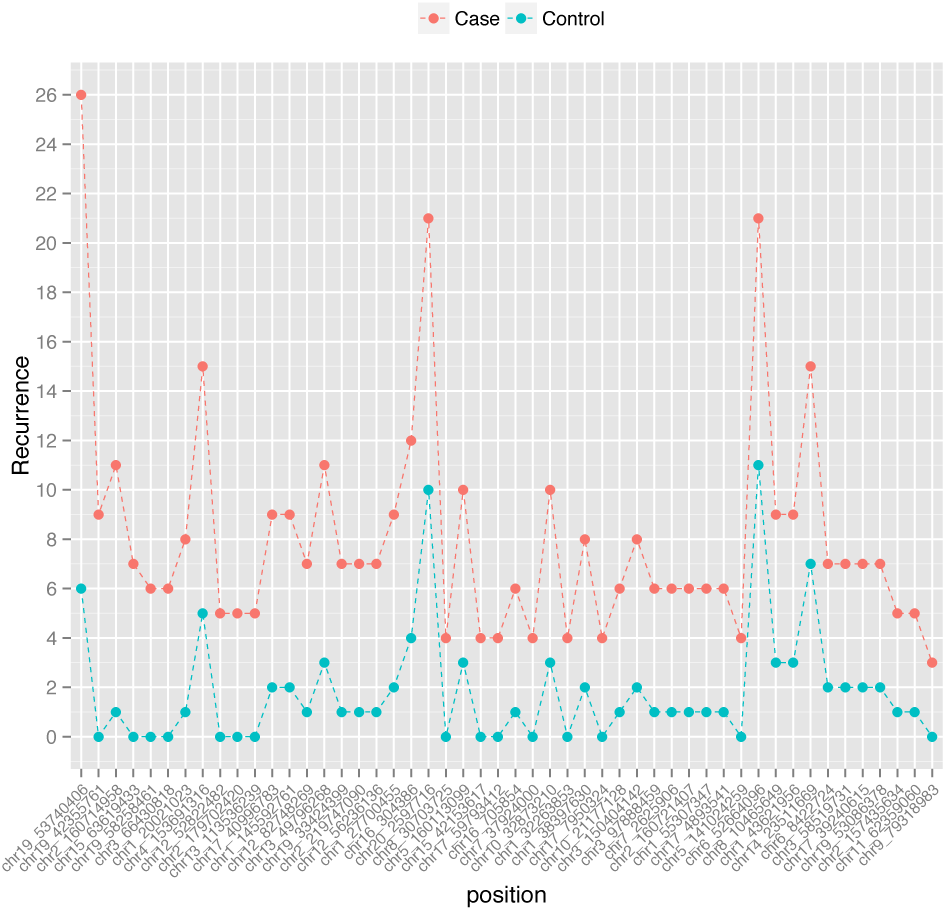
This plot shows the top 50 variants based on our method. The variants (horizontal axis) are ordered on decreasing order of the rank, meaning that the first variant is the top 1 and the last the top 50. The red dots represent the recurrence of the respective variant in cases, and the blue dots the recurrence in controls.

**Fig. 5.**
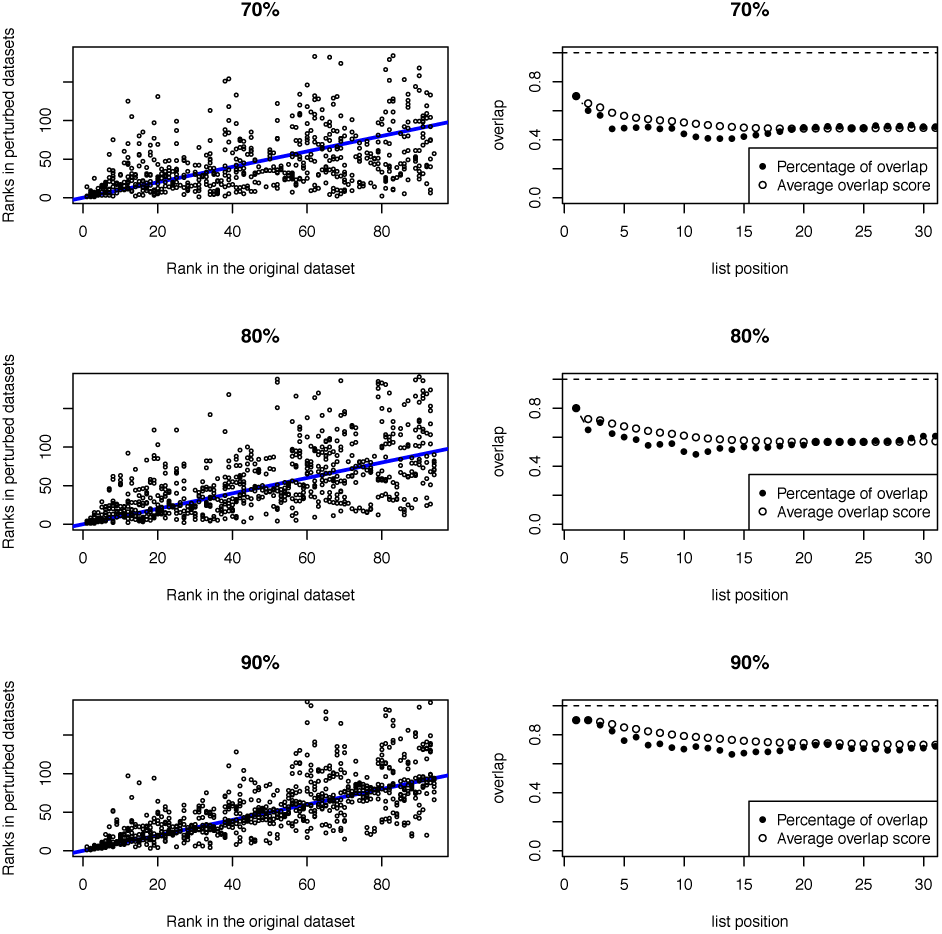
The plots on the left show the rank of all **10** iterations of the specific sample size proportion compared to the rank on the original dataset The blue line is presented as a guideline for comparison; it represents the case where the results of the algorithm in a smaller data provide the exact same ranking as the original. The plots on the right side show both the average percentage of overlap (white dots) and the average overlap score (black dots) of all 10 iterations over list position k.

As guidelines for analyzing the data based on our algorithm, we recommend further biological investigation of two main categories of genes. The first category consists of the genes at the top of the list where recurrence reflects the recurrence of a single variant within these genes. This means that the recurrence of a specific variant led the gene to be well ranked, even without the recurrence of other variants in the same gene. In the analyzed bipolar dataset, we observe the following genes in this category: ZNF677 (#2 gene level and #1 variant level); DMRTC2 (#3 in both gene and variant level); LRG1 (#11 gene level and #6 variant level). The second category consists of the genes that get highly ranked at the gene level due to overall recurrence of several variants within this gene. For example: ZNF776 (#4 gene and #10 variant); LY75/LY75-CD302 (#1 gene and #3 variant); ATP11A (#6 gene #12variant).

We checked our top results from this bipolar disorder study for known association to psychiatric disorders or brain expression. Supporting findings are summarized in table 1.

### 3.4 The robustness of the algorithm

We tested the robustness of the algorithm by evaluating the stability of the ranking when less data is available, meaning smaller sample size than the original. We showed good performance even with 70% of the original data. For this purpose, we randomly excluded some of the samples from the original data, and re-ran our algorithm on this dataset. To evaluate the similarities between the rankings of the different subsets to the original we use the R package GeneSelector (Boulesteix and Slawski, 2009). We applied the algorithm to nine different subsets of the original dataset ranging from only 10% to 90% of the original sample data. For each subset we generated ten different instances. GeneSelector was originally developed for testing stability and aggregation of ranked gene lists in gene expression data. Since all of the comparative metrics are based only on the ranking of the gene list, and not on the gene expression values, we could adapt it to our purpose. We focus on the following measures of evaluation: the percentage of overlap and the overlap score. The overlap of a fixed number *t* of top-variants is defined by the size *s*(*t*) of the intersection between two ranked lists i.e. the total number of variants that were ranked in the top-*t* of both lists. The overlap percentage is this overlap size, *s*(*t*), divided by *t*. For more details see (Boulesteix and Slawski, 2009). The second measure, the overlap score, is a weighted version of the overlap, where high ranked variants get higher weights than variants ranked below it. Specifically, we applied the linear weighted overlap score, meaning that position *i* in the list has a weight of 1/*i* on the score. As expected, as the sample size decreases compared to the original sample size, the intersection decreases as well. In this dataset, we can observe that when the sample size is 70% of the original size the algorithm provided results that the minimal average overlap score is 0.476, as seen in figure 5. Note that this is the average overlap score when it is computed based on the overlap of the top 19 variants. When the sample size is 80% the minimal average overlap score is 0.567, when considering the overlap of the top 21 variants. When the sample size is 90% the minimal average overlap score is 0.73, when considering the overlap of the top 31 variants. These show the robustness of our algorithm, that even when downsizing the data by 10%, the resulting ranking of the algorithm is relatively similar to the original ranking algorithm. Figures corresponding to datasets of sizes 10% till 60% of the original data are in supplementary figure 3 and 4.

**Fig. 5.**
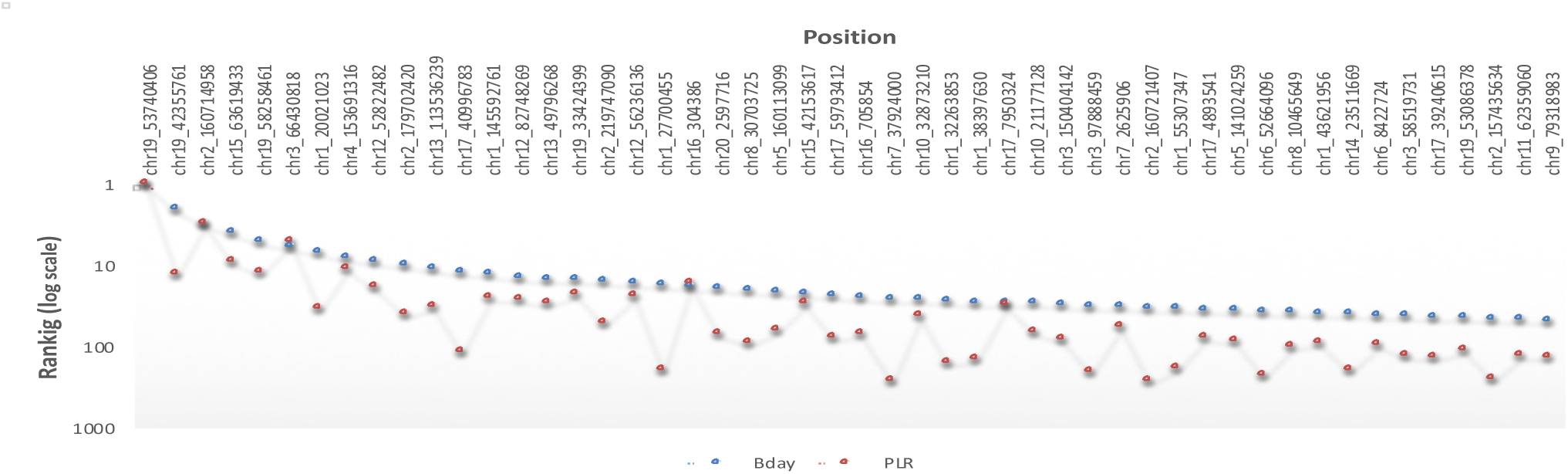
This plot shows the top 50 variants in the bipolar data based on our method and ordered by its rank of the variant level analysis. The red dots represent the rank of the respective variant based on PLR, and the blue dots the rank based on our algorithm.

## 4 Discussion

We have proposed to take advantage of a well-known probabilistic model–the Birthday Model- to indicate if findings in complex disorders may be real or are mere coincidence in the small number regimes expected for rare variant studies. Its analogy to our genomic problem allows an intuitive understanding of what we want to evaluate: whether the finding that we have observed is a true signal or a chance occurrence. The core probability of our method is determined by the observed recurrence, the sample size and the size of the gene. This input data for the algorithm can be easily extracted from the sequencing data. This is also especially beneficial in complex disorders where the genetic architecture is still unrevealed. Therefore, using a simplistic approach of observing coincidence of recurrence may help avoid bias due to inclusion of biological features that may not affect the causality of variants. Other biological criteria, such as type of mutation and minor allele frequency, can be included a priori, as a filtering step before implementing the algorithm.

A benchmark dataset for a complex disease with validated rare variants and known non-associated variants would be the ideal dataset for testing the ability of our algorithm to accurately select true causal variants. Unfortunately, there is no complex disease with a clear set of causative genes explaining a large fraction of the heritability. Therefore, we opted to explore the features of our method using simulations showing several rare variant scenarios detected by our method. We have also analyzed results of published studies in complex disorders under the implementation of the birthday decision-making method, showing the ability of the model to provide a quantitative metric to support the findings. And finally, we have performed the analysis of a case-control study on bipolar disorder using our algorithm, showing potentially interesting results for follow up studies. This same data on bipolar disorder was used to test the robustness of the algorithm, which showed good performance even when sub-sampled down to 70% of the original data. In addition to illustrating the robustness of the algorithm, this test shows the sensitivity of the algorithm when a smaller number of samples are available.

Here we have applied the birthday model for two different study designs of complex diseases: case-control and family studies looking for de novo mutations. However, the simplicity of the model gives the flexibility to adapt it to different studies, as for example in cancer for detection of mutations in tumor cells by modeling the normal cells as controls and the cancer cells as cases. It also can be applied for detecting protective variants by focusing on the recurrence of the variants present in the unaffected individuals instead the ones in the affected. This model can help to estimate the significance of the finding, especially for studies which are intended to identify rare mutations associated to complex disorders. The core probability based on the birthday model is very sensitive to the relatively low values of recurrence, which generally do not provide enough signal in existing statistical tests. With that, we may miss the moderate and common variants. However, existing tools showed good performance for these kinds of variants. In summary, given the insufficient sample size in current studies of complex disorders, our algorithm complements existing methods, specifically for the detection of the missing rare variants. Our approach provides a quantitative metric for evaluating whether rare findings, such as rare variants, may be meaningful in a world of coincidences. Hopefully, it will aid researchers in the prioritization of findings which merit further investigation.

## Acknowledgements

The authors thank Molly Gale Hammell for her comments on this work.

Exomes of the Synaptome project were sequenced at the CSHL Sequencing Shared Resource, supported by grant R01MH087992 **from** the National Institute of Health (NIH).

This work has been supported by a gift from T. and V. Stanley and the grant R01MH102068 from the National Institute of Health (NIH).

## References

Agarwala, V., et al. Evaluating empirical bounds on complex disease genetic architecture. Nature genetics 2013;45(12): 1418–1427.

Bigos, K.L., Haynes, R. M., Chen, D., Weinberger, D. R.. Genome-wide association study of olanzapine pharmacokinetics. In, 65th Annual Meeting of The American Society of Human Genetics. Baltimore, MD; 2015.

Boulesteix, A.L. and Slawski, M. Stability and aggregation of ranked gene lists. Briefings in bioinformatics 2009;10(5):556–568.

Boycott, K.M., et al. Rare-disease genetics in the era of next-generation sequencing : discovery to translation. Nat Rev Genet 2013;14(10):681–691.

Chong, J.X., et al. The Genetic Basis of Mendelian Phenotypes: Discoveries, Challenges, and Opportunities. American journal of human genetics 2015;97(2): 199–215.

Codina-Sola, M., et al. Integrated analysis of whole-exome sequencing and transcriptome profiling in males with autism spectrum disorders. Molecular autism 2015;6:21.

Diaconis, P. and Mosteller, F. Methods for Studying Coincidences. J Am Stat Assoc 1989;84(408):853–861.

Do, R., et al. Exome sequencing identifies rare LDLR and APOA5 alleles conferring risk for myocardial infarction. Nature 2015;518(7537): 102–106.

Eichler, E.E., et al. Missing heritability and strategies for finding the underlying causes of complex disease. Nat Rev Genet 2010;11(6):446–450.

Firth, D. Bias Reduction of Maximum-Likelihood-Estimates. Biometrika 1993;80(1):27–38.

Goes, F.S., et al. Exome Sequencing of Familial Bipolar Disorder. JAMA psychiatry 2016;73(6):590–597.

Goodwin, S., McPherson, J.D. and McCombie, W.R. Coming of age: ten years of next-generation sequencing technologies. Nat Rev Genet 2016;17(6):333–351.

Hindorff, L.A., Gillanders, E.M. and Manolio, T.A. Genetic architecture of cancer and other complex diseases: lessons learned and future directions. Carcinogenesis 2011;32(7):945–954.

Hu, X., et al. A survey of rare coding variants in candidate genes in schizophrenia by deep sequencing. Molecular psychiatry 2014;19(8):857–858.

Iossifov, I., et al. The contribution of de novo coding mutations to autism spectrum disorder. Nature 2014;515(7526):216–221.

Johansen, C.T., et al. Excess of rare variants in genes identified by genome-wide association study of hypertriglyceridemia. Nature genetics 2010;42(8):684–687.

Kiezun, A., et al. Exome sequencing and the genetic basis of complex traits. Nature genetics 2012;44(6):623–630.

Kryukov, G.V., et al. Power of deep, all-exon resequencing for discovery of human trait genes. Proceedings of the National Academy of Sciences of the United States of America 2009;106(10):3871–3876.

Maher, B. Personal genomes: The case of the missing heritability. Nature 2008;456(7218): 18–21.

Manolio, T.A., et al. Finding the missing heritability of complex diseases. Nature 2009;461(7265):747–753.

McClellan, J. and King, M.C. Genetic heterogeneity in human disease. Cell 2010;141(2):210–217.

Mckinney, E.H. Generalized Birthday Problem. Am Math Mon 1966;73(4P1):385–387.

Mitchell, K.J. What is complex about complex disorders? Genome Biol 2012;13(1):237.

Mitchell, K.J. and Porteous, D.J. Rethinking the genetic architecture of schizophrenia. Psychological medicine 2011;41(1): 19–32.

Moutsianas, L., et al. The power of gene-based rare variant methods to detect disease-associated variation and test hypotheses about complex disease. PLoS genetics 2015;11(4):e1005165.

Narasimhan, M., Bruce, T.O. and Masand, P. Review of olanzapine in the management of bipolar disorders. Neuropsychiatric disease and treatment 2007;3(5):579–587.

Poelmans, G., et al. Integrated genome-wide association study findings: identification of a neurodevelopmental network for attention deficit hyperactivity disorder. The American journal of psychiatry 2011;168(4):365–377.

Pritchard, J.K. and Cox, N.J. The allelic architecture of human disease genes: common disease-common variant…or not? Human molecular genetics 2002;11(20):2417–2423.

Rietkerk, T., et al. Network analysis of positional candidate genes of schizophrenia highlights myelin-related pathways. Molecular psychiatry 2009;14(4):353–355.

Rodriguez, J.M., et al. APPRIS: annotation of principal and alternative splice isoforms. Nucleic acids research 2013;41 (Database issue):D110–117.

Shendure, J., et al. DNA sequencing at 40: past, present and future. Nature 2017;550(7676):345–353.

Stranneheim, H. and Wedell, A. Exome and genome sequencing : a revolution for the discovery and diagnosis of monogenic disorders. J Intern Med 2016;279(1):3–15.

Strauss, K.A., et al. A population-based study of KCNH7 p.Arg394His and bipolar spectrum disorder. Human molecular genetics 2014;23(23):6395–6406.

Tennessen, J.A., et al. Evolution and functional impact of rare coding variation from deep sequencing of human exomes. Science 2012;337(6090):64–69.

Tomasik, J., et al. Pretreatment levels of the fatty acid handling proteins H-FABP and CD36 predict response to olanzapine in recent-onset schizophrenia patients. Brain, behavior, and immunity 2015.

Tomppo, L., et al. DISC1 conditioned GWAS for psychosis proneness in a large Finnish birth cohort. PloS one 2012;7(2):e30643.

van Bon, B.W., et al. CEP89 is required for mitochondrial metabolism and neuronal function in man and fly. Human molecular genetics 2013;22(15):3138–3151.

van der Zwaag, B., et al. Gene-network analysis identifies susceptibility genes related to glycobiology in autism. PLoS One 2009;4(5):e5324.

Westfall, P.H. and Young, S.S. Resampling-based multiple testing : examples and methods for P-value adjustment. New York: Wiley; 1993.

Wu, M.C., et al. Rare-variant association testing for sequencing data with the sequence kernel association test. American journal of human genetics 2011;89(1):82–93.

Yoshida, K., et al. Frequent pathway mutations of splicing machinery in myelodysplasia. Nature 2011;478(7367):64–69.

Zollner, S. Sampling strategies for rare variant tests in case-control studies. European journal of human genetics : EJHG 2012;20(10): 1085–1091.

Zuk, O., et al. Searching for missing heritability: designing rare variant association studies. Proc Natl Acad Sci U S A 2014;111(4):E455–464.

